# Hog1 acts in a Mec1-independent manner to counteract oxidative stress following telomerase inactivation

**DOI:** 10.1101/2023.10.04.560866

**Authors:** Bechara Zeinoun, Maria Teresa Teixeira, Aurélia Barascu

## Abstract

Replicative senescence is triggered when telomeres reach critically short length and activate permanent DNA damage checkpoint-dependent cell cycle arrest. Mitochondrial dysfunction and increase in oxidative stress are both features of replicative senescence in mammalian cells. Here, we show that reactive oxygen species (ROS) levels increase in the telomerase-negative cells of *Saccharomyces cerevisiae* during replicative senescence, and that this coincides with the activation of Hog1, a mammalian p38 mitogen-activated protein kinase (MAPK) ortholog. Hog1 activation is dependent on Pbs2, the MAPK kinase (MAPKK) in its canonical pathway, and counteracts increased ROS levels during replicative senescence. While Hog1 deletion accelerates replicative senescence, we found this could stem from decreased telomere length and reduced cell viability prior to telomerase inactivation. ROS levels also increase upon telomerase inactivation when Mec1, the yeast ortholog of ATR, is mutated, suggesting that oxidative stress is not simply a consequence of DNA damage checkpoint activation in budding yeast. We speculate that oxidative stress is a conserved hallmark of telomerase-negative eukaryote cells, and that its sources and consequences can be dissected in *S. cerevisiae*.

## Introduction

Telomeres are essential structures found at the ends of linear eukaryotic chromosomes, consisting of DNA sequences, proteins, and long non-coding RNA (LncRNA) telomeric transcripts (Shay & Wright, 2019). Telomeres crucially safeguard chromosome integrity by protecting against degradation and fusion events (Jain & Cooper, 2010). However, due to the “DNA end-replication problem”, telomeres gradually shorten with each cell cycle. Telomerase, a specialized reverse transcriptase, counteracts telomere shortening by adding repetitive telomeric sequences to chromosome ends. In human somatic cells, the telomere-protective functions become compromised when the telomeres shorten with cell divisions due to telomerase inactivation coupled with the “DNA end-replication problem”. When telomere lengths become critically short, they activate an irreversible DNA damage checkpoint-dependent cell cycle arrest, known as replicative senescence (Campisi & d’Adda, 2007; d’Adda *et al*, 2003). The unicellular eukaryote, *Saccharomyces cerevisiae*, relies on telomerase for its long-term viability (Lundblad & Szostak, 1989), but similar to human somatic cells, telomerase inactivation in budding yeast also leads to replicative senescence. When budding yeast cells divide in the absence of telomerase, they cease proliferation and enter a metabolically active state, arresting in the G2/M phase of the cell cycle (Enomoto *et al*, 2002; Ijpma & Greider, 2003). This cell cycle arrest in *S. cerevisiae*, which is akin to mammalian cells, relies on the activation of the DNA damage checkpoint kinases, Mec1 and Tel1 (the yeast orthologs of ATR and ATM, respectively), in addition to Rad53 phosphorylation (Abdallah *et al*, 2009; Khadaroo *et al*, 2009; Teixeira, 2013). Remarkably, not only is the triggering of replicative senescence in response to short telomeres evolutionarily conserved, but also many other essential telomeric functions and maintenance mechanisms (Kupiec, 2014; Wellinger & Zakian, 2012).

Studying replicative senescence is challenging due to the inherent heterogeneity resulting from intracellular differences in telomere lengths and the immense intercellular variations (Xu & Teixeira, 2019). Intriguingly, data collected from various organisms indicate that mitochondrial defects, oxidative stress, and chronic inflammation can accelerate telomere shortening and dysfunction (Ahmed & Lingner, 2018, 2020; Fouquerel *et al*, 2019; Opresko *et al*, 2002). These factors are potential sources of cell-to-cell variation and contribute to genome instability during replicative senescence (Passos & von Zglinicki, 2005). Notably, senescent human fibroblasts exhibit modifications in mitochondrial structure and function, accompanied by elevated reactive oxygen species (ROS) levels and oxidative damage (Hutter *et al*, 2004; Mai *et al*, 2010; Passos *et al*, 2010; Passos *et al*, 2007; Sitte *et al*, 2001; Sitte *et al*, 2000).

Similar metabolic alterations have also been observed in budding yeast during replicative senescence. A previous study revealed that the absence of telomerase resulted in increased mitochondrial mass, and a transcriptomic analysis indicated that energy production genes were up-regulated and stress response genes were induced (Nautiyal *et al*, 2002). However, data regarding ROS level alterations and their regulation during senescence in budding yeast is currently lacking.

P38, a member of the mitogen-activated protein kinase (MAPK) family, is critical for various cellular processes, including cellular senescence and oxidative stress responses (Martínez-Limón *et al*, 2020). In budding yeast, the MAPK Hog1, the ortholog of mammalian p38, is crucial for the defence against many stressors, including osmotic (Brewster *et al*, 1993) and oxidative stress (Haghnazari & Heyer, 2004). The canonical pathway of Hog1 activation involves two branches, the Sho1 and Sln1 branches, which converge to activate the MAPK kinase (MAPKK), Pbs2 (O’Rourke *et al*, 2002). Pbs2 then interacts with and phosphorylates Hog1 at the conserved residues, Thr^174^ and Tyr^176^, leading to its activation. Hog1 is a multifunctional protein with important functions in both the cytoplasm and nucleus, and it is vital for stress adaptation (de Nadal & Posas, 2022). Its roles encompass regulating gene expression by activating transcription factors, participating in gene initiation and elongation, regulating the cell cycle, and contributing to various steps in mRNA metabolism. Notably, Hog1 is activated in response to H_2_O_2_ stress (Haghnazari & Heyer, 2004) and regulates antioxidant genes by activating the transcription factors, Msn2/Msn4 (Wong *et al*, 2003) and Sko1 (Rep *et al*, 2001). In the absence of Hog1, cells become more sensitive to H_2_O_2_, which was previously shown to correlate with reduced expression of the *TSA2* gene (Wong *et al*., 2003). Conversely, sustained Hog1 activation can lead to cell death, which has been linked to alterations in mitochondrial respiration and increases in ROS levels (Vendrell & Posas, 2011). Hog1 counters this ROS increase by inducing *PNC1* and activating Sir2. Multiple studies have demonstrated that uncontrolled Hog1 activation disrupts mitochondrial function and elevates ROS levels, underscoring the critical importance of regulating Hog1 activation (Barbosa *et al*, 2012; Zhou *et al*, 2009). Furthermore, Hog1 has been implicated in autophagic processes and is required for mitophagy, the selective process of mitochondria degradation (Aoki *et al*, 2011; Huang *et al*, 2020; Mao *et al*, 2011). Interestingly, Hog1 also positively regulates the localization of the Sir complex to telomeres following osmotic stress and the silencing of telomeric regions (Mazor & Kupiec, 2009).

Here, we report that ROS levels increase during replicative senescence in budding yeast. During this process, Hog1 is activated by Pbs2 and plays a role in ROS detoxification. This countering of ROS increase occurs independently from the actions of Mec1. We also find that autophagy does not participate in replicative senescence in budding yeast. However, Hog1 participates in maintaining telomere length homeostasis and affects cell viability. Our results thus suggest that Hog1 serves as a link between telomeres and ROS metabolism.

## Results

### ROS levels increase during replicative senescence in budding yeast

Human senescent cells exhibit increased ROS levels during replicative senescence, however, data relating to *S. cerevisiae* senescent cells is lacking (Passos *et al*., 2010; Passos *et al*., 2007). Budding yeast telomerase is constitutively active but experimentally inactivating it triggers replicative senescence (Teixeira, 2013). We used a validated *TetO2-TLC1* system, where *TLC1*, which encodes the telomerase RNA template, is controlled by a repressible promotor by doxycycline, enabling the conditional shut-off of telomerase (Bah *et al*, 2011; Khadaroo *et al*., 2009; Soudet *et al*, 2014). To measure ROS levels, we used DCF, a molecule that can be directly oxidized by ROS and produce fluorescence in quantities reflecting ROS levels, which we can quantify by flow cytometry. Replicative senescence was detected from day three following culture with doxycycline as cell proliferation capacity decreased (Figure 1A). We also observed a simultaneous increase in ROS levels in the absence of telomerase (Figure 1B). These results were recapitulated in strains where *TLC1* was deleted; in these strains, we also found that ROS levels declined as cultures recovered their initial proliferation capacity following the emergence of post-senescence survivors (Figure S1). These data indicate that similar to mammalian models, budding yeast telomerase-negative cultures exhibit increased ROS levels.

**Figure 1:**
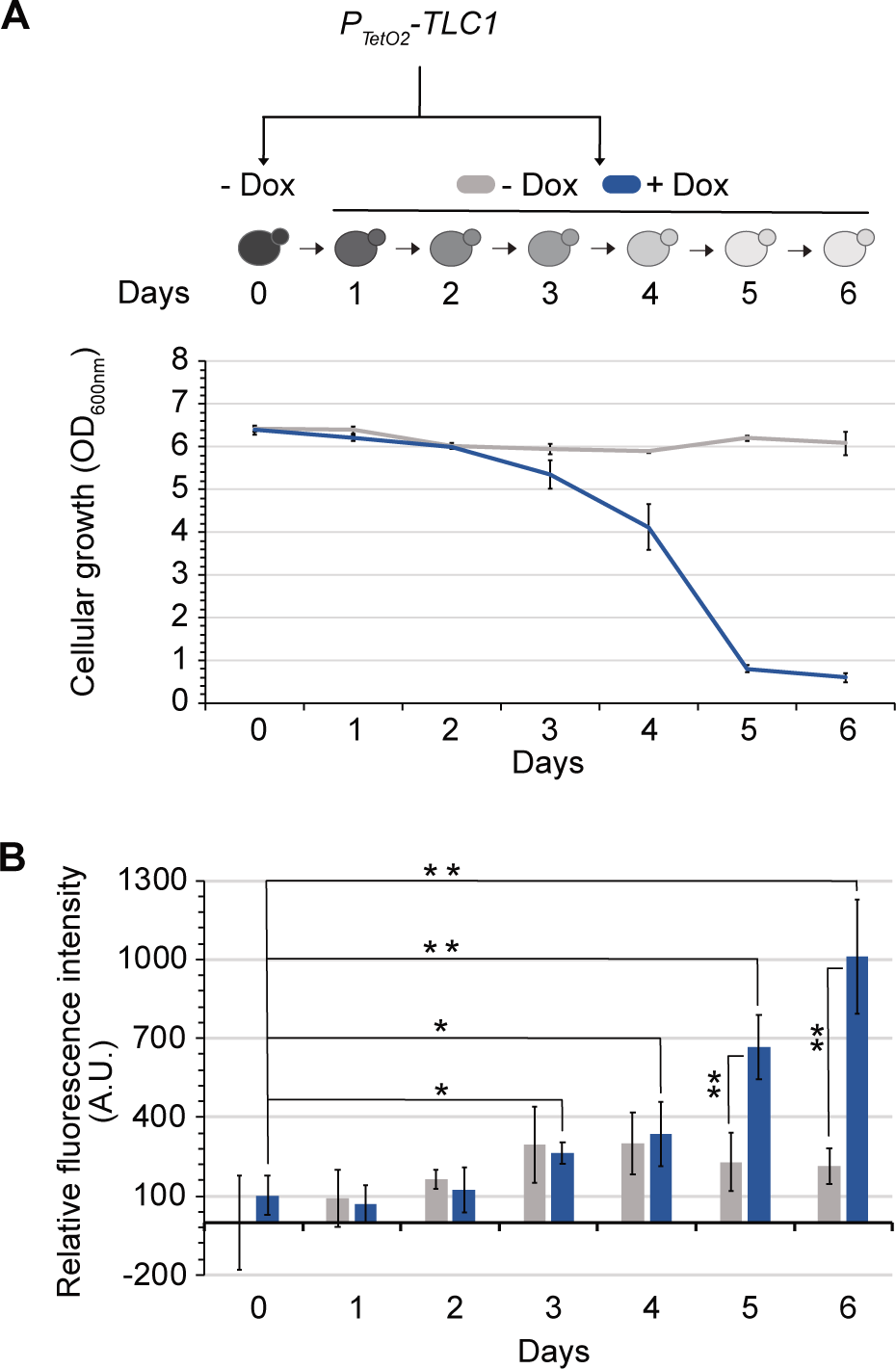
ROS levels increase during replicative senescence in budding yeast. Each consecutive day, cells with the genotype indicated were diluted in media either with or without doxycycline (Dox) to enable conditional shut-off of telomerase, and grown for 24 hours. Cell density at OD_600nm_ **(A)** and ROS levels **(B)** are plotted as mean ± SD of three independent clones. P-values were calculated by two-tailed Student’s t-test (* < 0,05; ** < 0,01) and only significant differences have been represented.

### Hog1 is activated during replicative senescence in a Pbs2-dependent manner and counteracts increase in ROS levels

We next investigated how ROS are regulated during replicative senescence. We focused on the multifunctional MAPK, Hog1, which is activated by oxidative stress (de Nadal & Posas, 2022; Haghnazari & Heyer, 2004). To determine whether Hog1 activation occurred during replicative senescence, we prepared protein extracts from senescent cultures and used a specific antibody to detect phosphorylated forms of Hog1. We observed that Hog1 was phosphorylated during replicative senescence from day three, concomitant with an increase in ROS levels (Figure 2 C and B, respectively and S2). This phosphorylation was dependent on Pbs2, the MAPKK that precedes Hog1 in its canonical pathway (O’Rourke *et al*., 2002). We also observed that *HOG1* or *PBS2* deletion resulted in a similar premature loss of cell viability, when telomerase was inactivated (Figure 2 A). This suggests that the Hog1 pathway may play a role in senescent cells. While the HOG1 pathway activates transcription factors of antioxidant genes to reduce ROS levels (Vendrell & Posas, 2011), excessive Hog1 activity can increase ROS levels by disrupting mitochondrial respiration (Barbosa *et al*., 2012; Vendrell & Posas, 2011). To understand which of these Hog1 functions was involved in replicative senescence, we measured ROS levels in the absence or presence of Hog1 during senescence. Our results showed that the increases in ROS levels started earlier and reached higher levels in *hog1Δ* compared to *HOG1* strains (Figure 2B). This suggests that increases in ROS levels trigger an oxidative stress response that activates the Hog1 pathway, which is required for ROS detoxification in telomerase-inactivated cells.

**Figure 2:**
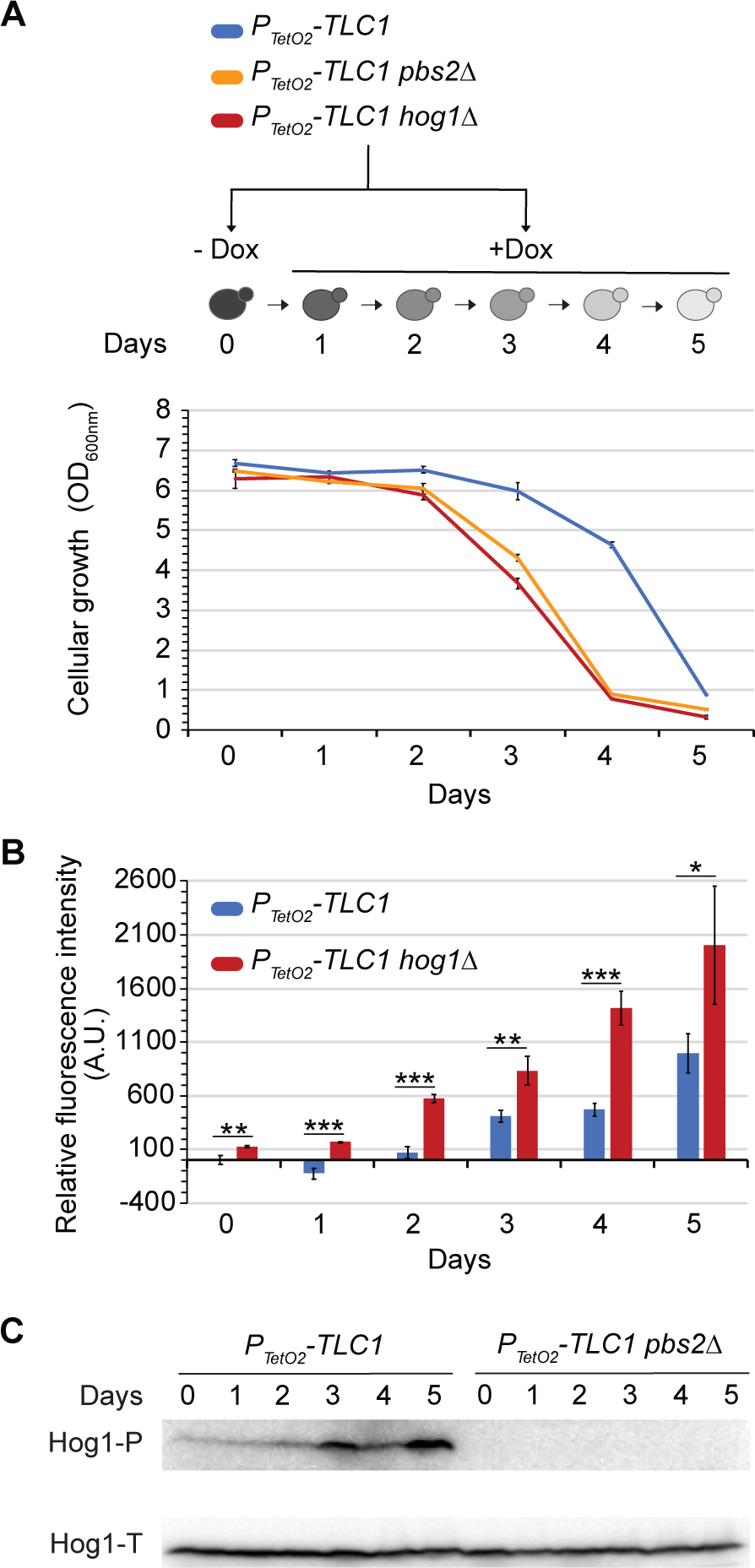
Hog1 is activated during replicative senescence in a Pbs2-dependent manner and counteracts increase in ROS levels. Cells with the genotypes indicated were treated as described as mean ± SD of three independent clones. P-values were calculated by two-tailed Student’s t-test (* < 0,05; ** < 0,01; *** < 0,001) and only significant differences between the two strains used have been represented. **(C)** Protein extracts analysed by Western blot using an antibody against phosphorylated forms of Hog1’s human ortholog p38 (Hog1-P), or total Hog1 (Hog1-T).

### *HOG1* deletion affects telomere length homeostasis and cell viability

Given the direct relationship between replicative senescence and telomere shortening, we investigated the potential influence of Hog1 on telomere length homeostasis prior to telomerase inactivation, and the rate of telomere shortening in the absence of telomerase. We thus performed telomere-PCR on DNA samples from *HOG1* and *hog1Δ* strains to determine telomere length. Our results showed that *HOG1* deletion resulted in slightly shorter telomeres of ∼30 bp prior to telomerase inactivation (Figure 3A-B, S3A). However, no significant differences in telomere shortening rates were observed in the absence of telomerase in either the *HOG1* or *hog11* strains; shortening rates were measured to be approximately 2.5 bp/cell population doubling (Figure 3C-D, S3B), similar to previously published results (Marcand *et al*, 1999; Soudet *et al*., 2014). We therefore conclude that Hog1 contributes to the maintenance of telomere length homeostasis. As telomere length homeostasis results from a balance between telomere lengthening by telomerase and telomere shortening due to the “DNA end-replication problem”, we suggest that Hog1 might interfere with telomerase recruitment or activity.

**Figure 3:**
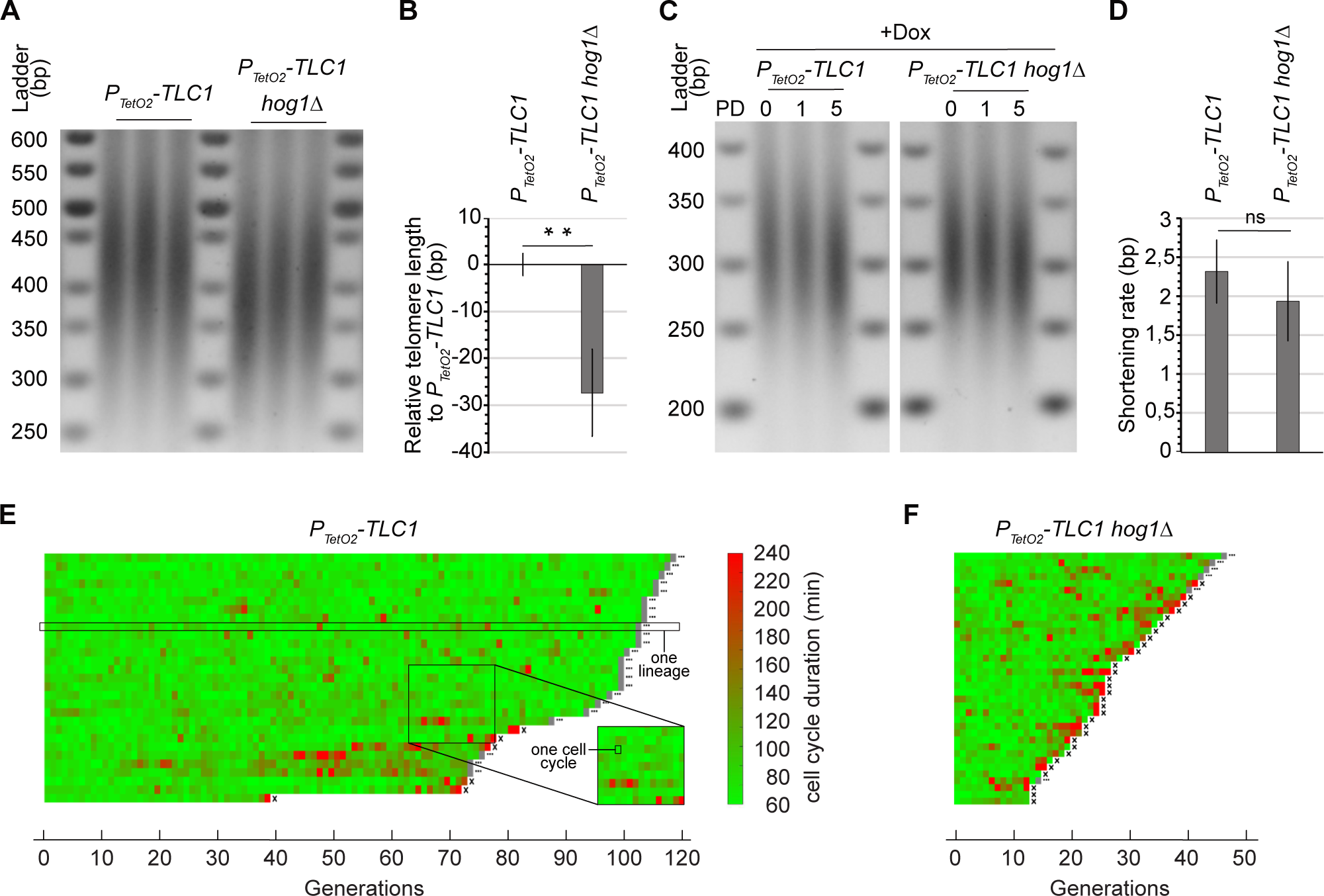
*HOG1* deletion affects telomere length homeostasis and cell viability. Representative telomere-PCR of Y’ telomeres from the strains indicated **(A)** and their quantification, plotted as mean ± SD **(B)**. P-values were calculated by two-tailed Student’s t-test (** < 0,01). **(C)** Cells of the genotypes indicated were pre-cultured overnight in doxycycline-containing media to enable telomerase shut-off. Cells were then diluted and grown in the same media for the indicated population doublings (PD). Representative telomere-PCR of Y’ telomeres from genomic DNA extracts are shown. **(D)** Quantification of telomere shortening was measured between PD 1 to 5 and plotted as mean ± SD. **(E,F)** Microfluidics results of independent lineages with the genotypes indicated. Cells were introduced into the microfluidics microcavities and cultured in SD. Each horizontal line represents the consecutive cell cycles (generations) of a single lineage, and each segment corresponds to one cell cycle. An ellipsis (…) at the end of the lineage line indicates that the cell was living after the experiment, whereas an X indicates cell death. Cell cycle duration is indicated by the coloured bar.

We reasoned that the accelerated senescence we observed in the *hog1Δ* strains when telomerase was inactivated could be due to shorter telomere lengths prior to telomerase inactivation, rather than because *HOG1* is required for cell growth in the absence of telomerase. To confirm this, we employed a microfluidics system, which allows consecutive cell cycles from individual cell lineages (herein referred to as lineages) to be tracked, thereby enabling more precise quantification of cell proliferation (Xu *et al*, 2015). In the presence of telomerase, *HOG1* cells grew indefinitely with a spontaneous mortality rate of ∼0,38% (Figure 3E). In contrast, the absence of *HOG1* caused the mortality rate to increase to ∼5,8% even in the presence of telomerase (Figure 3F). This has not previously been observed with cell population growth (liquid or solid). This data indicates that *HOG1* loss alone causes some cell death, which, when combined with telomerase inactivation, could have contributed to the accelerated senescence we observed.

Collectively, these findings suggest that the apparent acceleration of senescence we observed in the absence of Hog1 could be attributable to a marked increase in intrinsic mortality rates combined with the initial shorter telomeric lengths. However, this does not preclude a significant role for Hog1 in detoxifying ROS during replicative senescence, particularly as *hog1Δ* strains exhibit much higher ROS levels throughout replicative senescence compared to *HOG1* strains (Figure 2B).

### Autophagic processes do not modify the onset of replicative senescence

Autophagy is a process that involves self-eating and bulk degradation, where organelles and their components are delivered to vacuoles to be degraded and recycled (Yin *et al*, 2016). Autophagy can be selective, when specific cargo, such as damaged organelles, are degraded (Suzuki, 2013). The specific degradation of dysfunctional mitochondria is known as mitophagy. Hog1 plays a role in autophagy under certain conditions (Babele *et al*, 2018; Bicknell *et al*, 2010; Prick *et al*, 2006) and is considered to be a mitophagy activator (Aoki *et al*., 2011; Mao *et al*., 2011; Shen *et al*, 2017). We therefore investigated whether autophagic processes were essential during replicative senescence. We deleted the *ATG8* and *ATG32* genes, which encode two proteins essential for bulk autophagy and mitophagy in budding yeast, respectively (Suzuki, 2013). These mutants *atg8Δ* and *atg32Δ* displayed a blockage of bulk autophagy and mitophagy respectively, confirmed by an assay based on Rosella, a fluorescence-based pH biosensor (Rosado *et al*, 2008) (Figure S4).

We then investigated whether blocking these processes in wild-type cells would affect senescence dynamics. Liquid senescence assays revealed that senescence remained unchanged in the absence of autophagy or mitophagy, suggesting that these processes were not essential for the viability of telomerase-negative budding yeast cells (Figure 4A). Similarly, following telomerase inactivation, the senescence profiles remained unchanged when either *ATG8* or *ATG32* were deleted in a *hog1Δ* background (Figure 4A). We concluded that Hog1 activity occurs independently from autophagic processes, which do not alter the onset of replicative senescence in budding yeast.

**Figure 4:**
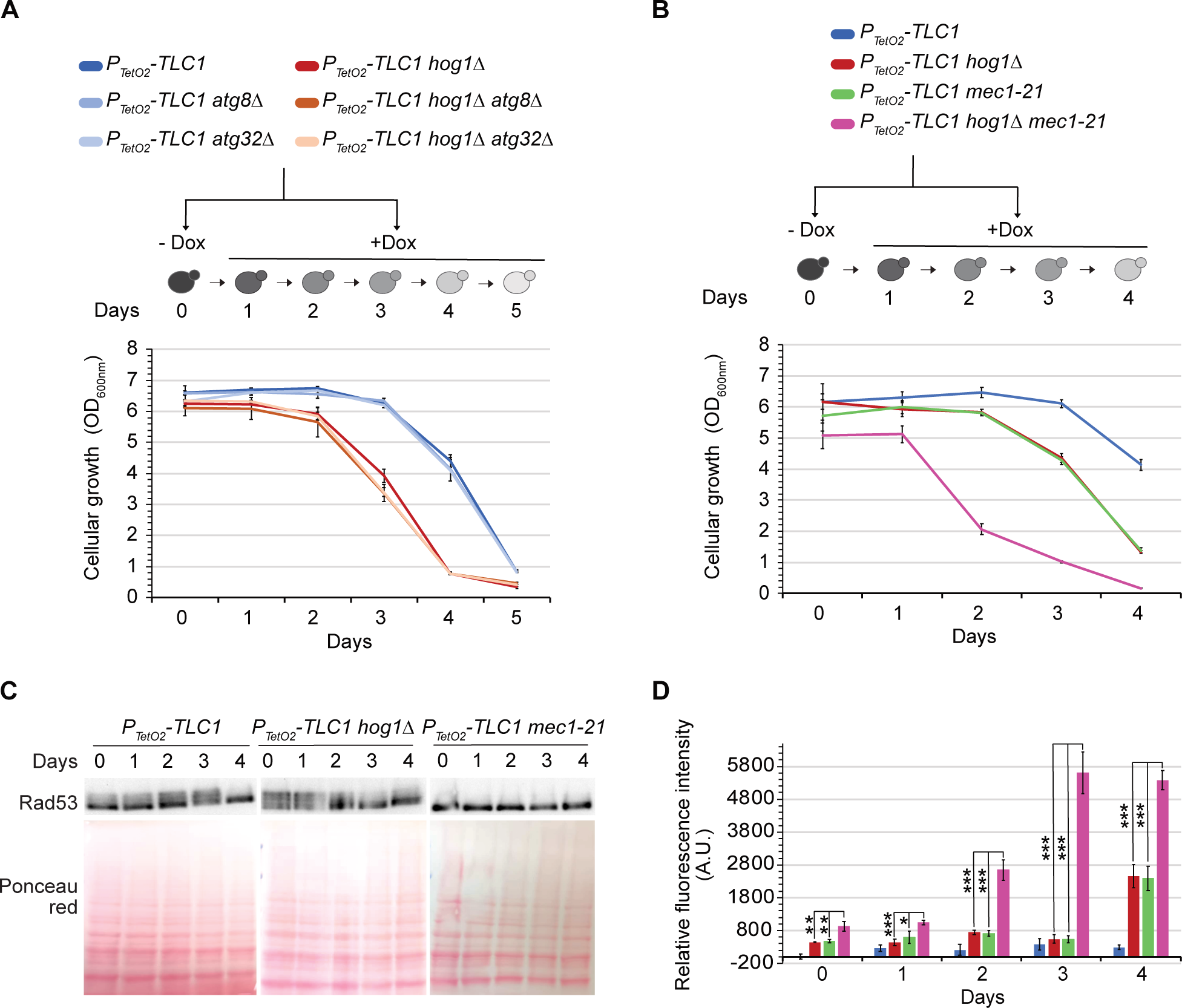
Hog1 interaction with multiple pathways. Cells with the genotypes indicated were treated as described in Figure 1. Cell density at OD_600nm_ **(A, B)** and ROS levels **(D)** are plotted as mean ± SD of three independent clones. P-values were calculated by two-tailed Student’s t-test (* < 0,05; ** < 0,01; *** < 0,001) and only significant differences between the triple mutant and the double mutants have been represented. **(B)** Representative telomere-PCR of Y’ telomeres from the strains indicated. **(C)** Protein extracts analysed by Western blot using an antibody against Rad53. The mobility shift of the band indicates Rad53 phosphorylation.

### Hog1 acts in a Mec1-independent manner to regulate ROS levels during replicative senescence

We hypothesized that ROS level increases could result from cells being in a senescent state. Mec1 is a pivotal kinase in budding yeast, essential for the DNA damage checkpoint and the onset of replicative senescence (Hector *et al*, 2012). Mec1 and the DNA damage checkpoint pathway are also known to protect cells against oxidative stress (Tsang *et al*, 2014). Hog1 and Mec1 are both necessary to combat the oxidative stress induced by hydrogen peroxide, but act independently (Haghnazari & Heyer, 2004). A hypomorph mutant of *MEC1, mec1-21,* contains a G to A substitution at position 2644, outside the kinase domain (Ritchie *et al*, 1999). The *mec1-21* mutant displays lower dNTP levels and shorter telomeres (∼50 bp) compared to wild type strains (Fasullo *et al*, 2009; Ritchie *et al*., 1999). The *mec1-21* mutant retains essential functions but is defective for the S phase checkpoint and Rad53 activation following UV and HU exposure (Fasullo & Sun, 2008; Sun & Fasullo, 2007). We thus used the *mec1-21* mutant to investigate whether the actions of Hog1 against ROS were Mec1-dependent during replicative senescence. We inactivated telomerase and measured cell growth and proliferation capacity over time in the P*_TetO2_-TLC1*, P*_TetO2_-TLC1 hog1Δ*, P*_TetO2_-TLC1 mec1-21*, and triple mutant strains. When compared to *MEC1* cells, we observed that *mec1-21* accelerated the loss of viability under telomerase-negative conditions (Figure 4B), consistent with the initial shorter telomeres. We also verified that Rad53 phosphorylation was impaired in the *mec1-21* strains (Figure 4C). Yet, under these conditions, where the DNA damage checkpoint was disabled, telomerase inactivation resulted in more pronounced increases in ROS levels. This indicates that Mec1 also participates in ROS detoxification in the absence of telomerase (Figure 4D). In addition, while the triple mutant, P*_TetO2_-TLC1 hog1Δ mec1-21*, displayed a much lower proliferation capacity, it showed an even more pronounced increase in ROS compared to the respective single mutants (Figure 4D). These results are consistent with a model where Hog1 and Mec1 are both involved in ROS detoxification during replicative senescence but act in independent pathways.

## Discussion

Here, we have shown that increased ROS levels are a feature of replicative senescence in budding yeast. Hog1, one of the five MAPKs of *S. cerevisiae*, is activated by Pbs2 during replicative senescence and counteracts increases in ROS levels, likely independently of Mec1. In addition, Hog1 regulates telomere length homeostasis, and its deletion results in a marked increase in cell mortality rates. Our findings also indicate that autophagic processes are not essential in the context of replicative senescence in budding yeast.

Previous studies have shown that Hog1 is activated in response to exogenous acute stresses, such as H_2_O_2_ exposure, where it is essential for triggering antioxidant genes and maintaining cell viability (Haghnazari & Heyer, 2004; Wong *et al*., 2003). Given our findings that ROS levels increase during replicative senescence, it is plausible that oxidative stress directly triggers Hog1 pathway activation. Replicative senescence is an endogenous process resulting from telomerase inhibition, that leads to numerous cellular modifications at both genomic and metabolic levels. Consequently, other modifications may also contribute to Hog1 activation. Notably, a previous study proposed that Hog1 activation in response to H_2_O_2_ stress primarily occurs through the Sln1-Ypd1-Ssk1-Ssk2-Pbs2 pathway, with Ssk2 acting as the MAPKKK that specifically activates Hog1 in response to oxidative stress, but not Sho1 branch (Lee *et al*, 2017). Therefore, determining the Hog1 pathway upstream of Pbs2 might help clarify the origin of Hog1 pathway activation in the absence of telomerase.

Microfluidics analysis, where cells grow individually, showed that *HOG1* deletion in the presence of telomerase increased cell mortality rates by ∼15 fold. This increase may have gone undetected in other studies where cells were grown in populations as colonies or liquid cultures due to competition and selection of the fittest cells. Similar mortality rates have been described for other mutants considered “viable”, underscoring the high sensitivity of the microfluidics method (Xu *et al*., 2015). We speculate that the Hog1 pathway might be important in response to certain intrinsic stresses, and that it becomes essential in rare situations. Accordingly, a potential role for Hog1 under normal stress-free cellular conditions, unrelated to telomeres, has been described (Reynolds *et al*, 1998).

We have shown that Hog1 participates in telomere length homeostasis in budding yeast. This could be due to Hog1 acting to positively regulate telomere transcriptional silencing through the localization of the Sir complex following osmotic stress (Mazor & Kupiec, 2009). It could also be that the absence of Hog1 disrupts subtelomeric heterochromatin, which would alter telomere length homeostasis.

In conclusion, this study has shown that the metabolic alterations observed in human senescent cells are conserved in budding yeast. These alterations involve a conserved MAPK Hog1/p38 pathway, although the outcome might differ in different species. As most basic functions in telomere biology are conserved in eukaryotes, determining the mechanistic link between telomere shortening and increases in ROS levels in budding yeast will be essential to clarify how telomeres have evolved in the context of eukaryotic evolution.

## Material and Methods

### Yeast strains

All yeast strains used in this study were derived from a W303 background corrected for *RAD5* and *ADE2* (Table 1). Gene deletions were constructed as previously described (Longtine *et al*, 1998). *Mec1-21* point mutation was constructed using Crispr-Cas9, as previously described (Lemos *et al*, 2018). Strains expressing Rosella constructs from plasmids were constructed as previously described (Rosado *et al*., 2008). Primers used are listed in Table 2.

**Table 1:**
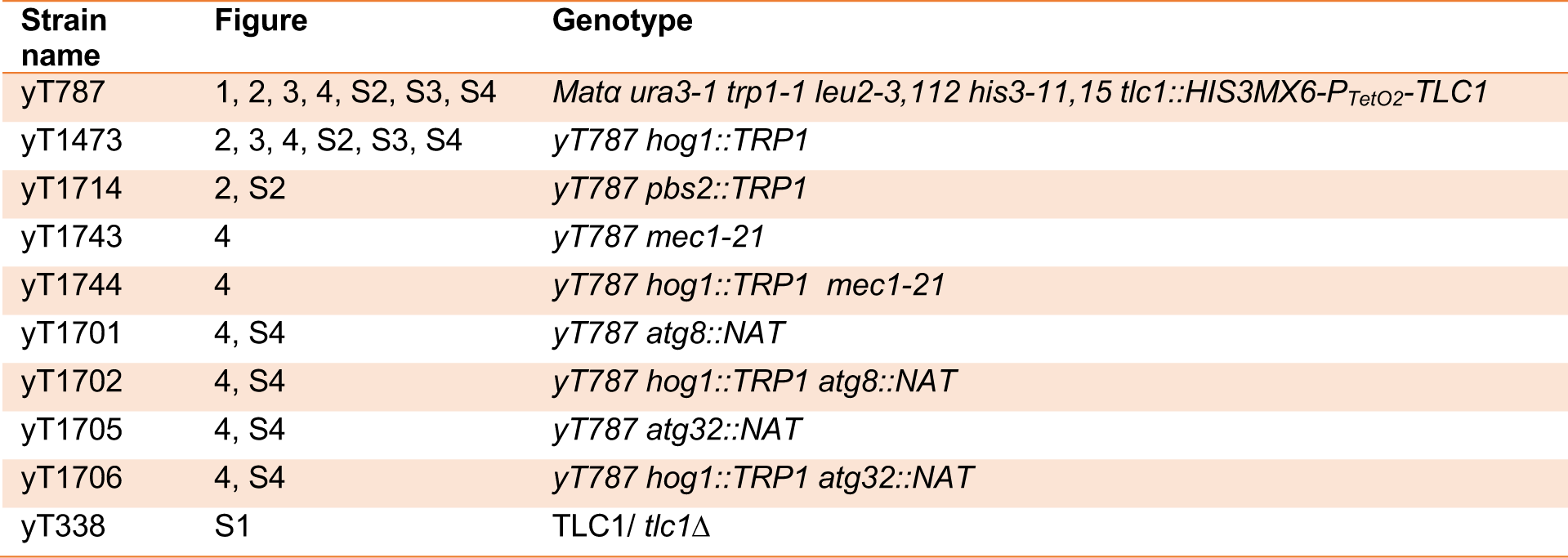
strains used in this study.

**Table 2:**
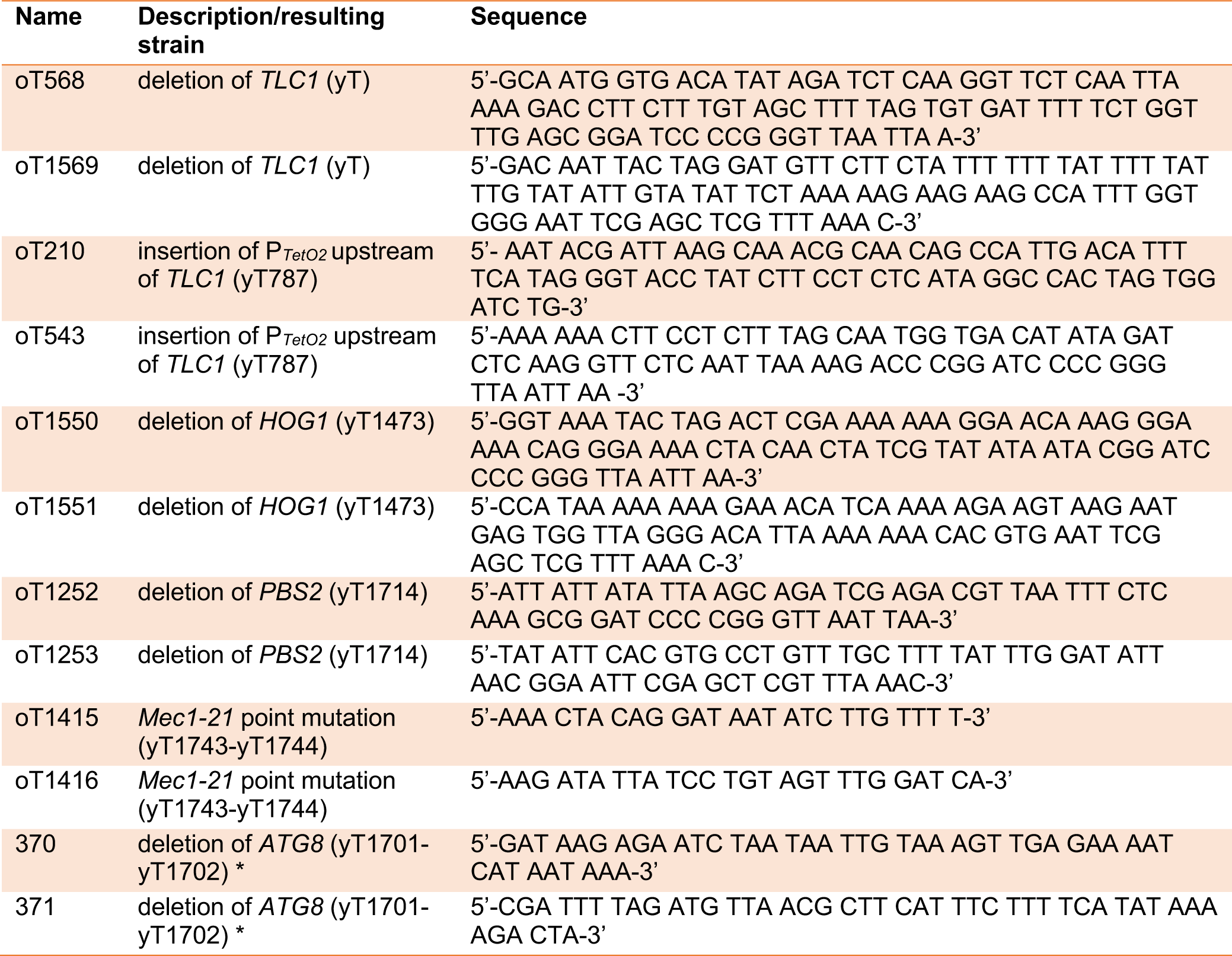
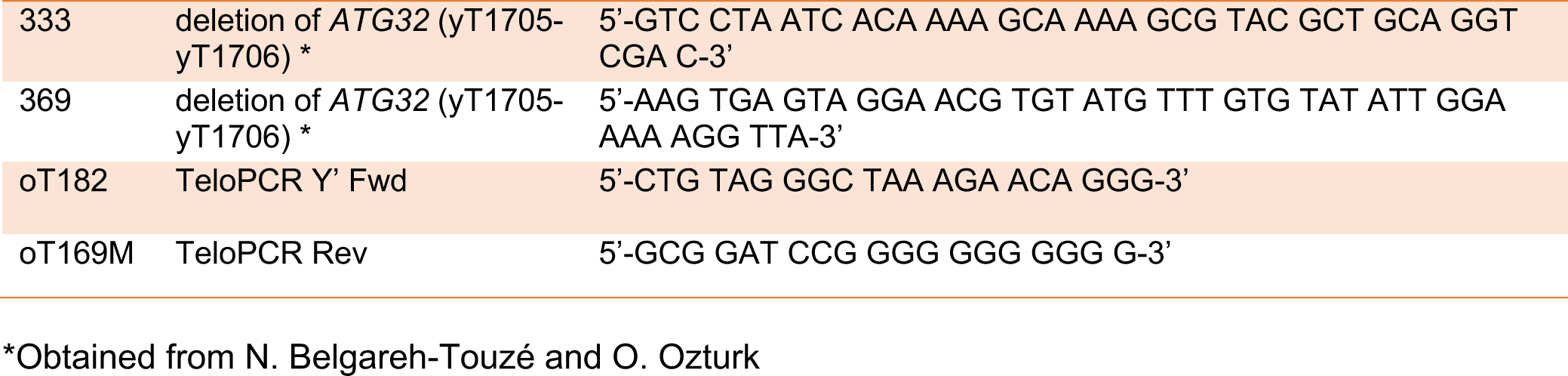
Primers used in this study.

### Liquid senescence experiments

Strains were grown at 30°C in liquid-rich media (YPD). Cell suspensions were diluted to 0,001 OD_600nm_ with a final concentration of 30 µg/mL of doxycycline *(Sigma-Aldrich #D9891),* and the OD_600nm_ was measured after 24 hours. Cultures were similarly diluted for several days and daily samples were taken for analysis.

### ROS detection

Yeast cultures with an OD_600nm_ of 0,4 were incubated at 30°C in darkness for one hour in 500 μL of sterile 1X PBS, containing DCF-DA (2’,7’-Dichlorofluorescin diacetate) *(Sigma-Aldrich #D6883)* at a final concentration of 10 μM. Samples were washed and then resuspended in 500 μL of 1X PBS. Fluorescence was then analysed by flow cytometry using the settings, 488 (λex)/533 (λem) in Accuri C6 or MACSQuant Analyzer 10. The mean intensity values were then plotted. The mean values of *P_TetO2_-TLC1* at day 0 without doxycycline were subtracted.

### DNA extraction

Cells with an OD_600nm_ of 5 were centrifuged for four minutes at 2000 g and washed in 500 μL of sterile distilled water. After centrifugation, 200 μL of lysis buffer (Triton 100 X - 10% SDS sodium dodecyl sulfate - 5 M NaCl – 0,5 M EDTA ethylenediaminetetraacetic acid - 1 M Tris - H2O), 200 μL of 0,45 μm acid-washed glass beads *(Sigma-Aldrich #G8772)*, and 200 μL of phenol: chloroform: isoamyl alcohol solution *(25:24:1, Sigma-Aldrich #77617)* were added to the cell pellet. The tubes were vortexed for 15 minutes at 4°C, followed by the addition of 200 μL of TE (Tris/EDTA) at pH 8. After five minutes of centrifugation at maximum speed, the aqueous phase was transferred to a new tube containing the same volume of isopropanol. After mixing by inversion, the samples were centrifuged for one minute at maximum speed. The resulting DNA pellet was washed in 500 μL of ethanol. Finally, after centrifuging and drying in a speed vacuum (three minutes at 40°C), the DNA pellet was resuspended in 50 μL of TE and 0,1 μL of RNase A (100 mg/ml) and incubated for 30 minutes at 37°C. The quality and integrity of the DNA were checked by agarose gel electrophoresis (1% agarose, 0,5X TBE). The quantity was evaluated using Qubit2 *(Thermofisher)*. Samples were stored at −20°C.

### SDS-PAGE and Western blot

Cells with an OD_600nm_ of 5 were collected by centrifugation. The pellet was lysed in 0,2 M of sodium hydroxide (NaOH) for 10 minutes on ice. After adding trichloroacetic acid (TCA) at a final concentration of 0,5%, the samples were incubated again on ice for 10 minutes. After centrifuging for 10 minutes at maximum speed at 4°C, the pellet was resuspended in 50 μL of Laemmli 4X buffer and denatured at 95°C for five minutes. Protein samples were electrophoresed on a 10% denaturing gel (or 7,5% for Rad53 detection) of 37,5:1 Acrylamide:Bis-acrylamide *(Sigma-Aldrich #A3699)*. Proteins were then transferred to a nitrocellulose membrane *(Amersham Protran 0.45 NC, GE HealthCare)* and stained with Ponceau red. The following antibodies were used: *Cell Signalling, #9211* to detect Phospho-Hog1, Santa Cruz, *#sc-165978* to detect total Hog1, *Abcam, #ab104252* to detect both unphosphorylated and phosphorylated forms of Rad53, and the horseradish peroxidase-coupled secondary antibody (HRP) *(Sigma, #A9044 and #A9169)*. The signals were revealed using an electrochemiluminescence reagent *(ClarityWestern ECL, Biorad)* and recorded using the ChemiDoc Imaging System (Biorad).

### Telomere-PCR

This method was adapted from (Forstemann *et al*, 2000). In brief, 40 ng of genomic DNA was denatured at 98°C for five minutes before the tailing reaction in 20 μL of *New England Biolabs* Buffer 4 (1X), 100 μM of dCTP, and 1U of terminal transferase *(New England Biolabs #M0315L)*. The reaction was incubated at 37°C for five minutes, followed by five minutes at 94°C, and then maintained at 4°C. For the PCR reactions, 5 μL of the polyC-tailed products were used with 1X Taq Mg-free Buffer *(New England Biolabs)*, 500 nM of each primer (Table 2), 200 μM of dNTPs, and 1 U of Taq Standard Polymerase (*New England Biolabs #M0273*) in a final volume of 30 μL. The following PCR program was used: three minutes at 94°C, followed by 45 cycles of 20 seconds at 94°C, 15 seconds at 57°C, 20 seconds at 72°C, and finally, five minutes at 72°C. The TeloPCR products were then loaded onto a large 2% agarose gel with 1X TBE buffer and 100 ng/ml of ethidium bromide (BET). A 50bp molecular weight marker was also loaded *(New England Biolabs #N3236)*. Electrophoresis was then performed at 50V for 15 hours. Visualization and analysis were performed using the ImageLab® software *(Biorad)*.

### Microfluidics analysis

Microfluidics analyses were performed as previously described (Xu *et al*., 2015).

## Acknowledgements

We wish to thank Naima belgareh-Touzé and Oznür Ozturk for sharing reagents and technical advice, the Teixeira lab, and the UMR8226 unit members for technical support and fruitful discussions. We also wish to thank Pascale Jolivet, Prisca Berardi, Yann Lustig, Juan Manuel Peralta, Pol Ubeda, Victoria Rojat and Clara Basto for technical help, and Francesc Posas’ lab, Miguel Godinho Ferreira, and Zhou Xu for fruitful discussions. Work in MTT’s laboratory was supported by the CNRS, Sorbonne University, the “Fondation de la Recherche Medicale” (“Equipe FRM EQU202003010428”), by the French National Research Agency (ANR) as part of the “Investissements d’Avenir” Program (LabEx Dynamo ANR-11-LABX-0011-01) and The French National Cancer Institute (INCa_15192). BZ. is a recipient of fellowship from the «Ministère de l’enseignement supérieur et de la Recherche» (MESR).

---

**Supplementary Figure 1:**
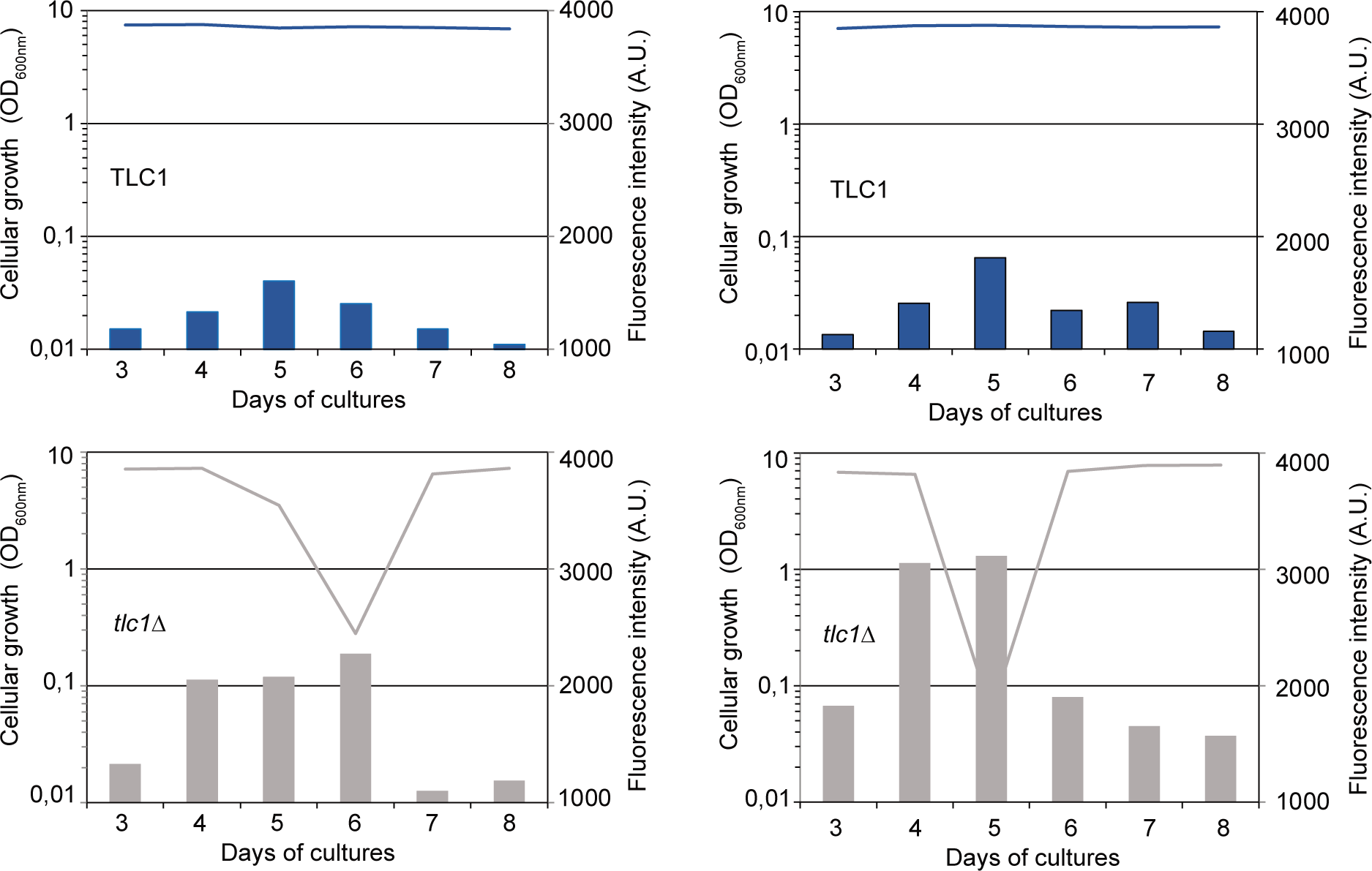
ROS levels increase during replicative senescence in *tlc1Δ* strains. A diploid *TLC1/tlc1Δ* strain was dissected, and the four spore-derived colonies obtained after two days were pre-cultured. Each consecutive day, cultures were diluted, as described in Figure 1, and grown for 24h. Cell density at OD_600nm_ (curve-left axis) and ROS levels (histogram-right axis) are plotted.

**Supplementary Figure 2:**
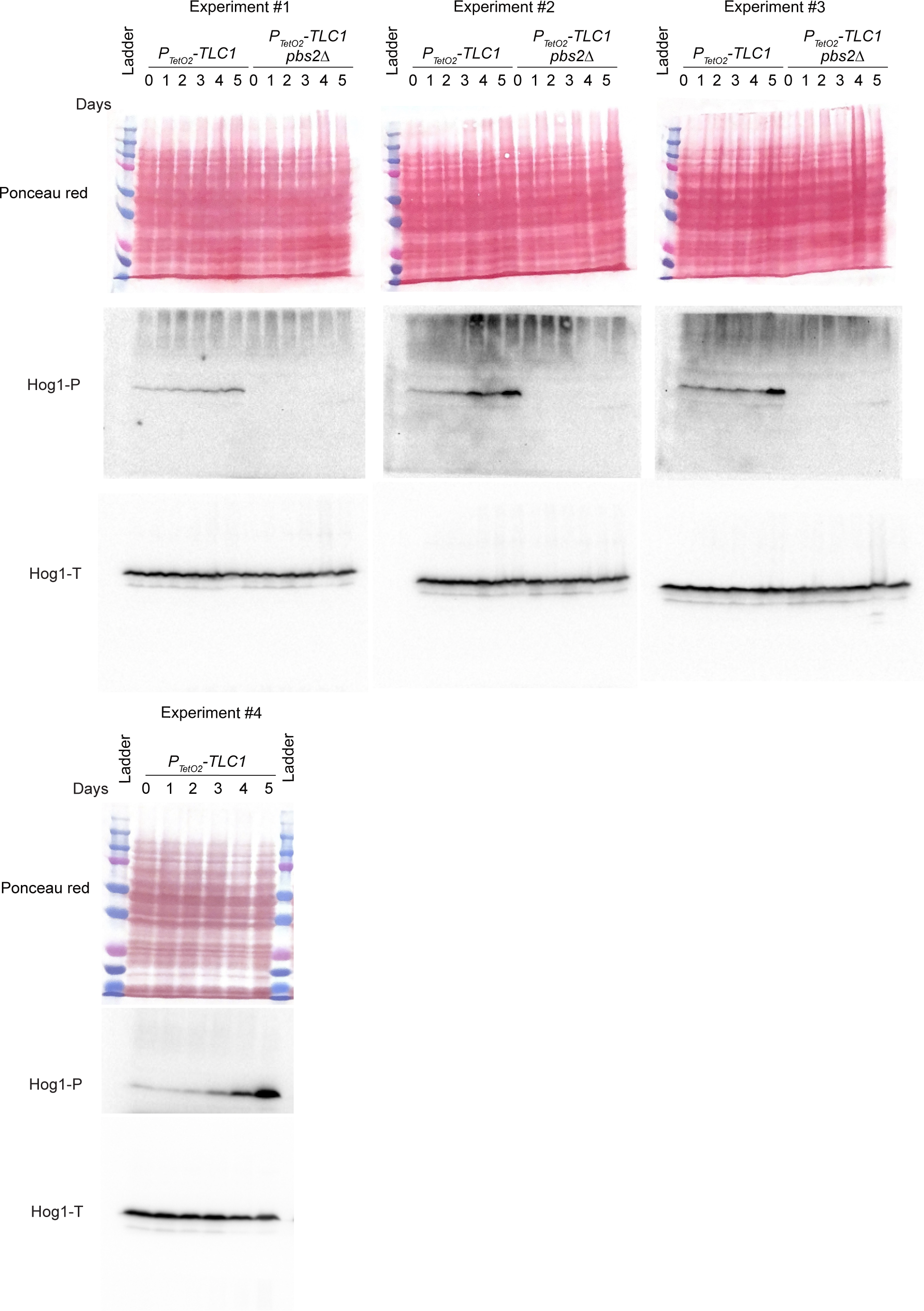
Western blot of 4 independent experiments relative to Figure 2. Protein extracts analysed by Western blot using an antibody against the phosphorylated forms of Hog1 human ortholog p38 (Hog1-P), or total Hog1 (Hog1-T).

**Supplementary Figure 3:**
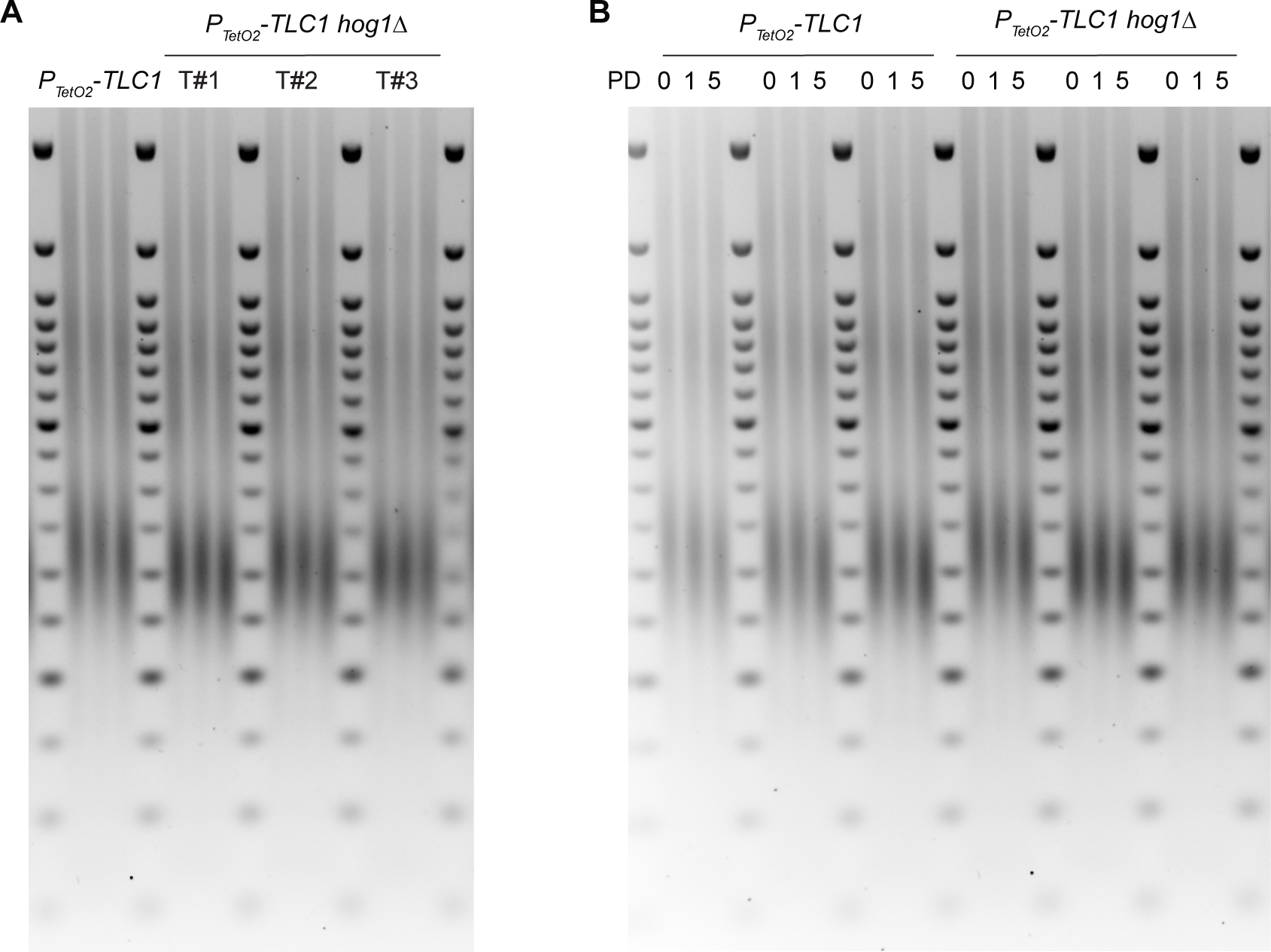
*HOG1* deletion affects telomere length homeostasis. **(A)** Technical triplicates of telomere-PCR of Y’ telomeres from the independent transformants (T#1-3) and strains indicated. **(B)** Technical triplicates of the experiment described in Figure 3C. Population doublings (PD).

**Supplementary Figure 4:**
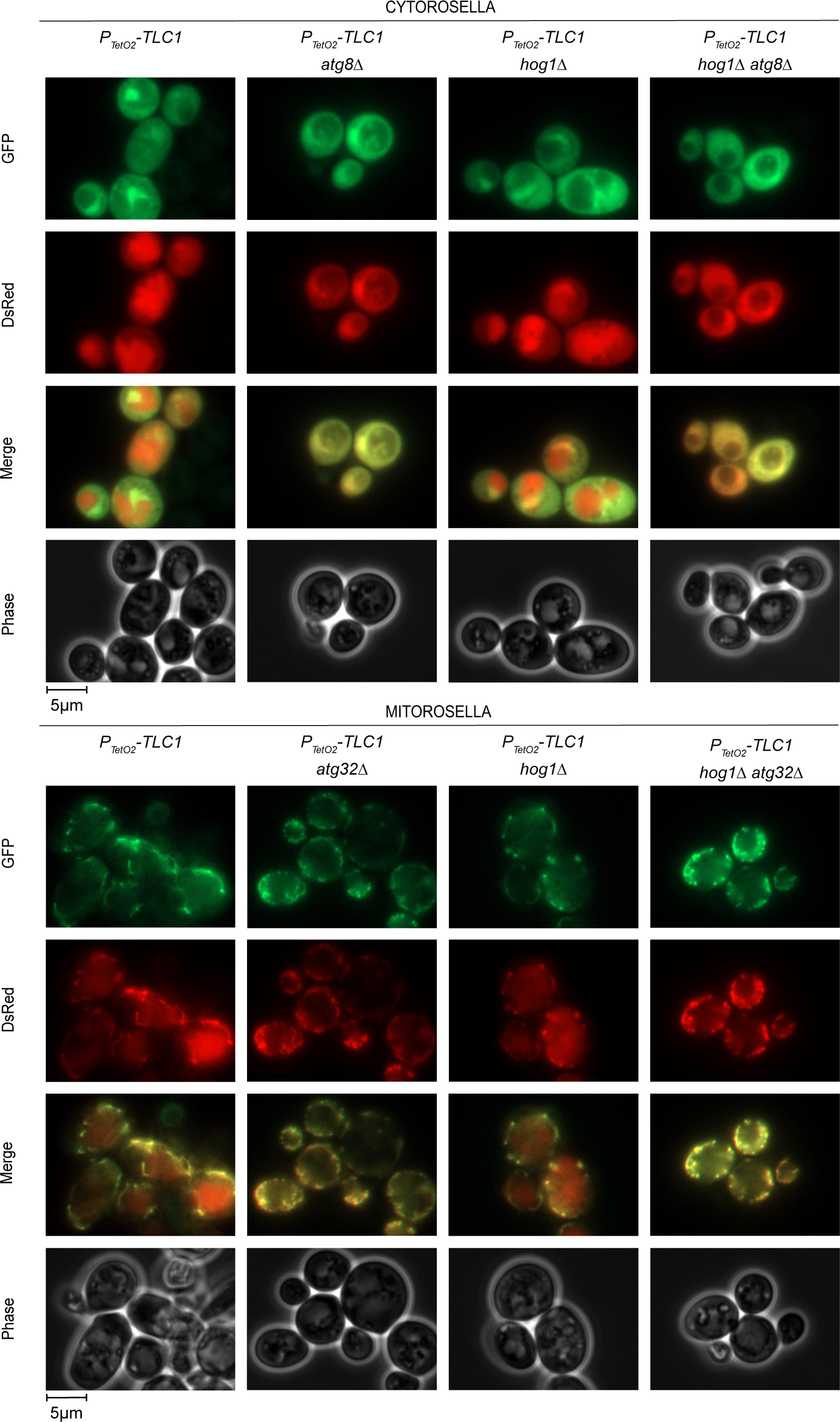
Strains deleted for *ATG8* and *ATG32* show an autophagy- and mitophagy-impaired phenotype, respectively. Images of fluorescence microscopy following autophagy and mitophagy induction by nitrogen depletion for 24 hours. Cytorosella is a fusion between a cytoplasmic targeting signal, a DsRed, and a GFP sensitive to pH. Mitorosella is similar to cytorosella but with a mitochondrial targeting signal.

